# Combined administration of novobiocin and apomorphine mitigates cholera toxin mediated cellular toxicity

**DOI:** 10.1101/2022.12.13.520354

**Authors:** Sonali Eknath Bhalerao, Himanshu Sen, Saumya Raychaudhuri

## Abstract

Cholera is a dreadful disease. The scourge of this deadly disease is still evident in the developing world. Though several therapeutic strategies are in practice to combat and contain the disease, there is still a need for new drugs to control the disease safely and effectively. Keeping in view the concern, we screened a small molecule library against a yeast model of cholera toxin A subunit. Our effort resulted in the discovery of a small molecule, apomorphine effective in reducing the lethality of toxic subunit in yeast model. In addition, novobiocin, an inhibitor of ADP ribosylation process, a key biochemical event through which cholera toxin exerts its action on host, was also found to rescue yeast cells from cholera toxin A subunit mediated toxicity. Finally, both molecules prevented cholera toxin mediated cellular toxicity on HT29 intestinal epithelial cells. We also observed that combined administration of both drug molecules worked better than single drug in countering toxin driven lethality.

## Introduction

*Vibrio cholerae* causes cholera, a life threatening diarrheal disease with worldwide distribution^1^. Out of 220 serogroups, strains belonging to serogroups O1 and O139 are associated with epidemic cholera. Though decades of continuing efforts have harvested wealth of information on the biology of the bacterium and disease per se, cholera continues to rage and remains a major public health problem in the developing and war torn countries. As per WHO records, over 3 million people are affected annually, leading to over 100,000 deaths^2^. Among multiple lines of treatments, oral or intravenous rehydration remains the mainstay in cholera. Antibiotics are also given as an adjunct therapy to control and decrease duration of the disease. But indiscriminate use of antibiotic increases in the emergence of multidrug resistant strains of *V. cholerae*^3,4^, thereby contributing towards global burden of AMR. Vaccines with limited efficacy providing short term protection are also available and can be useful to restrict the spread of cholera in outbreak prone area. But the vaccines can give only 65% protection against cholera for 6 months up to 3 years^5^. Therefore, there is a desperate need to come up with newer strategies to combat cholera.

With continuing effort from researchers across the globe, new areas are increasingly emerging to tackle the pathogenesis and fitness of the organism. So far, this is achieved either by chemically synthesized small molecules jeopardizing the function of regulatory proteins (e.g ToxT, LuxO) involved in the production of virulence factors^6–12^ or gut commensal derived metabolites targeting the overall fitness and survival of the organism in the host intestine^13–20^. Though these strategies hold a promise in controlling the disease, the safety and effectiveness of each strategy under clinical settings yet to be evaluated. Interestingly, natural variants of regulatory proteins are also identified and found resistant to small molecule mediated functional modulation^21^. Probiotic mediated septicemia is also evidenced in animal model as well as in ICU patients^22,23^ and raising grave concern over the usage of probiotic strains. Keeping view of all recent advancements and issues associated with different strategies to combat cholera, we were driven by a desire to explore further in the same direction with a hope to come up with better molecule for abating the disease.

*Saccharomyces cerevisiae*, the budding yeast has been exploited as a non-mammalian model system for functional evaluation of several virulence factors from many pathogenic bacteria including *Vibrio cholerae*^24–31^. Simple ectopic expression of virulence factors can lead to a spectrum of discernible phenotypes in yeast that aid to build testable hypotheses regarding their function and roles in pathogenesis. Not only advancing functional understanding of microbial and higher eukaryotic proteins, such yeast model systems are also instrumental in the discovery of novel small molecule modulators against specific proteins further prompted us to embark on the present work^32,33^.

In this work, we developed a yeast model of cholera toxin by ectopically expressing gene encoding cholera toxin A (CTXA) subunit. The expression of CTXA resulted a severe growth defect in the recombinant yeast. Subsequently, the CTXA expressing yeast was subjected to screen against small molecule library and one small molecule, apomorphine, was found to inhibit CTXA mediated growth retardation. In addition, we also observed another small molecule novobiocin, a known inhibitor of ADP ribosylation also reduced CTXA toxicity in recombinant yeast strain.

Cholera toxin is demonstrated to alter cellular morphology in PC12 cells^34^. We also documented morphological changes of HT29 intestinal cell lines mediated by commercially available cholera toxin (CT) and such cellular alterations was also reduced by exogenous administration of apomorphine and novobiocin. A combination of apomorphine and novobiocin exerted a stronger inhibitory effect on CT mediated lethality and promoted cellular survival under experimental conditions. Collectively, this study reveals the effectiveness of apomorphine, a dopamine agonist commonly used to treat Parkinson disease, in controlling cholera toxin mediated cellular toxicity and also demonstrates the strength of yeast model of virulence factors in drug discovery.

## Results

### Ectopic Expression of cholera toxin A subunit in *Saccharomyces cerevisiae* affects cell growth

The gene fragment containing cholera toxin A subunit (*ctxA*) without secretion signal was amplified and cloned under the control of GAL-1 promoter in high copy number vector pESC Leu to generate recombinant plasmid pESC-Leu-CTXA. To investigate the toxicity of CTXA, the recombinant plasmid pESC-Leu-CTXA was transformed into *S. cerevisiae* BY4741. Under inducing condition in the presence of galactose, spotting on solid agar media exhibited strong growth inhibition (**Figure 1A**). There was also weak growth inhibition under repressive (glucose) condition. This is due to leaky expression from GAL1 promoter of pESCLeu. The leaky expression issue related to GAL1/10 promoter has been noticed in previous studies as well^30^. The growth inhibition was recapitulated in the liquid growth assay (**Figure 1B**). It is documented that lethality of some effector proteins is also linked with the high level of expression^35^. In some cases, high level of expression also leads to non-specific toxic effects. To ascertain, low levels of expression of CTXA maintains similar toxicity, the gene was cloned in pGML10, a low copy number vector (Table 1). *S. cerevisiae* strain BY4741 transformed with pGML10-CTXA exhibited lethal phenotype in inducing condition (**Figure 1C**). In addition, gene encoding CTXA was also chromosomally integrated at HO-locus. Upon induction with galactose, the recombinant strain (BY4741/HOΔ::HO-CTXA-kanMX4-HO) exhibited CTXA mediated lethality (**Figure 1D**). It should be noted that remaining experiments were carried out in this recombinant strain harbouring chromosomally integrated singly copy of gene encoding CTXA.

**Figure 1.**
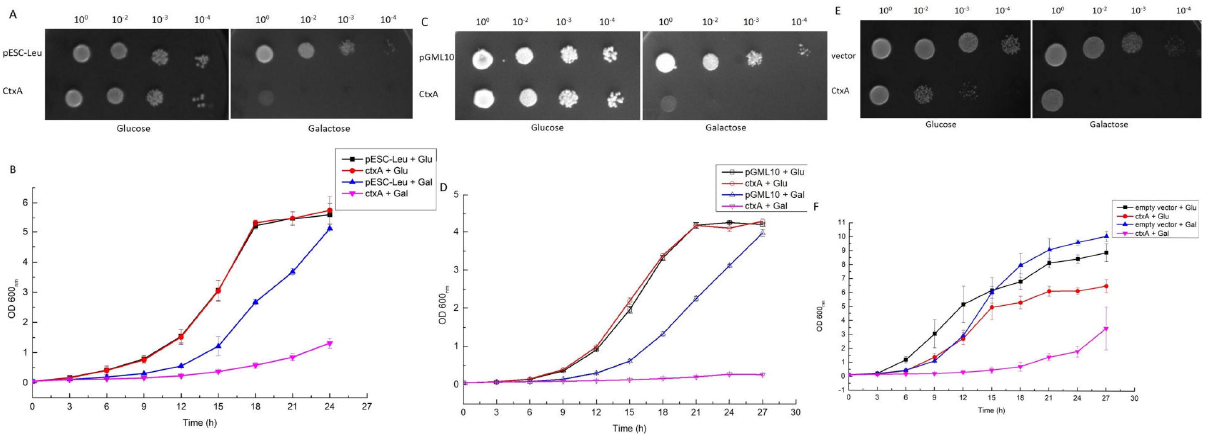
Effect of *ctxA* expression on yeast growth: **A and B. Effect of *ctxA* expression on yeast growth when expressed in high copy number vector:** *ctxA* was cloned in a high copy number vector pESC-Leu and was expressed in BY4741 strain of yeast. For liquid growth assay, the secondary cultures were set up at a starting OD_600_ 0.05 and were monitored for 24 hrs. OD_600_ was checked every 3 hrs. The data shown here is collected from 3 independent experiments. For spotting data, secondary cultures were set up at starting OD_600_ 0.1, after 6 hrs, cultures were serially diluted (10^0^, 10^2^, 10^3^, 10^4^) and were spotted on glucose and galactose plates. Pictures were taken after 60-70 hrs. **C and D. Effect of *ctxA* expression on yeast growth when expressed in a low copy number vector:** *ctxA* was cloned in a low copy number vector pGML10 and was expressed in BY4741 strain of yeast. Liquid growth assay and spotting assay was performed for 27 hrs as mentioned in **A. E and F. Effect of *ctxA* expression on yeast growth when single copy of gene is expressed:** *ctxA* was expressed in a chromosomal DNA of BY4741 using HO-pGAL-polykanMX4-HO vector. Liquid growth was performed for 27 hrs as explained above. For spotting experiment, the secondary cultures were grown to OD_600_ 0.8-0.9 and spotted on solid agar containing glucose and galactose.

### A small molecule apomorphine mitigated CTXA mediated growth retardation in recombinant yeast model system: Lessons from phenotypic screen

Thus far, our results suggested that expression of cholera toxin A subunit is detrimental for yeast growth. Therefore, any molecule perturbs the function of the CTXA should restore the growth of our recombinant yeast strain. Keeping this in mind, we engaged on a phenotypic screening assay to find small molecule modulator (s) of cholera toxin. To pursuit our interest, we subjected our recombinant *S. cerevisiae* strain harbouring *ctxA* gene (**Table 1**) to screen against small molecule library obtained commercially (e.g., Spectrum limited). The plate screening resulted one positive candidate molecule which appeared to rescue yeast viability (**Figure 2A**). The molecule is apomorphine, a dopamine agonist^36^. To evaluate further the efficacy of apomorphine in growth restoration of the recombinant yeast, liquid growth assay coupled with viability spotting was done with various concentrations of the drug. We observed that apomorphine showed better rescue of the recombinant strain if added twice in the cultures during the entire span of liquid growth assay (**Figure 2B, Figure 2C and Figure 2D**). Taken together, our study clearly demonstrated the utility of yeast model system in facilitating discovery of novel inhibitor against cholera toxin.

**Figure 2.**
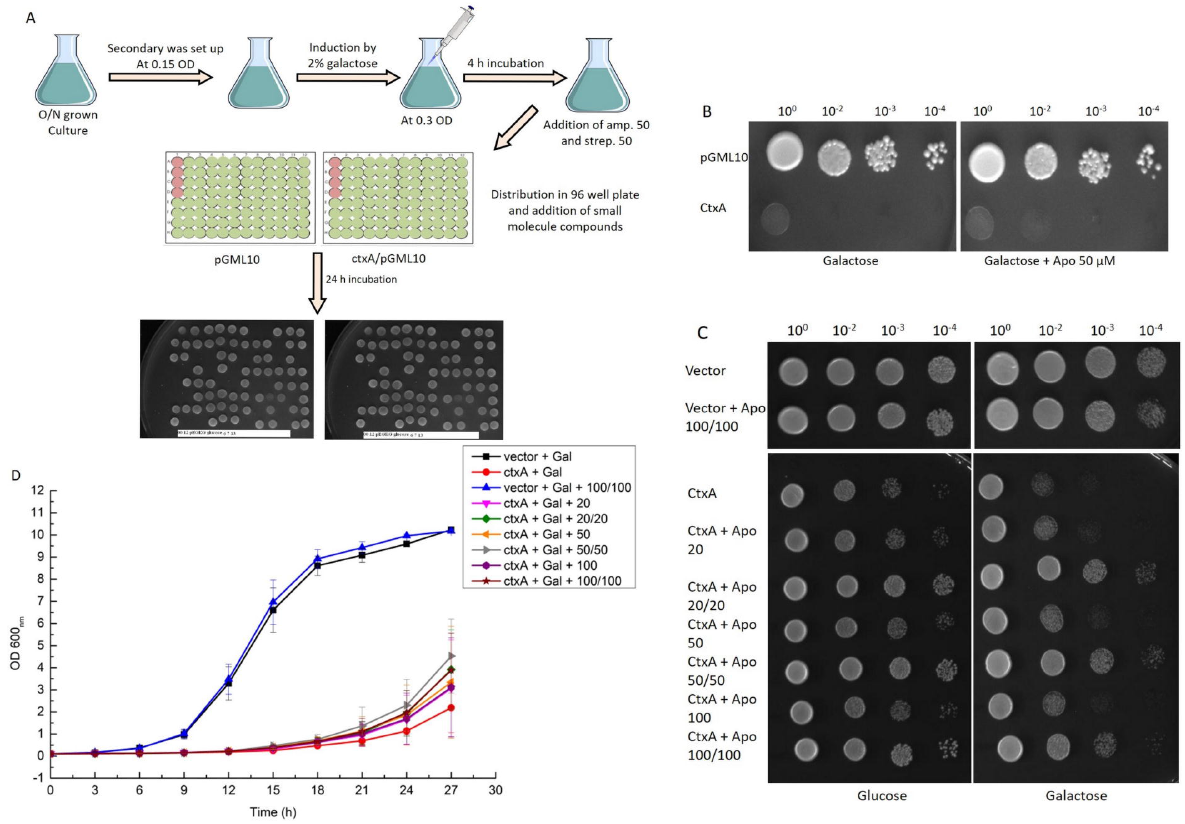
Small molecule library screening using cholera toxin yeast model system: **A. Schematic of high-through put screening:** Pre-induced cultures of BY4741/*ctxA*/pGML10 and BY4741/pGML10 as a control were used. These cultures were evenly distributed in 96 well plates, then compounds from the library plate were added at the final concentrations of 20 μM. Both plates were incubated for 24 hrs at 30 □ and were then spotted on glucose and galactose plates for readouts. **B. Tertiary screening of Apomorphine HCl:** Pre-induced cultures of BY4741/*ctxA*/pGML10 and BY4741/pGML10 as a control were spotted on solid agar plate containing galactose and galactose with apomophine (50 μM). Pictures were taken after 60-70 hrs **C and D. Effect of Apomorphine HCl on cholera toxicity in yeast:** Secondary cultures of BY4741/*ctxA*-HO and vector control were set up at starting OD_600_ 0.05 in glucose and galactose media with different concentrations of apomorphine and were monitored for 27 hrs. OD_600_ was checked every 3 hrs. The data shown here is collected from 3 independent experiments **(C)**. At the end point of liquid growth assay, the cultures were serially diluted (10^0^, 10^2^, 10^3^, 10^4^) and were spotted on glucose and galactose plates. Pictures were taken after 60-70 hrs **(D)**. Liquid growth assay.

### Novobiocin, an ADP ribosylation inhibitor rescued recombinant yeast from cholera toxin mediated lethality

Mechanistically, cholera toxin more specifically the enzymatic A subunit of cholera toxin activates Gsα by irreversible transfer of ADP moiety of NAD^+^ to Arg^201^ of Gsα^37–39^. The ADP-ribosylation of G protein triggers cascade of events involving hyper activation of cellular adenylate cyclase, followed by a sharp rise in intracellular level of cyclic AMP. The increase c-AMP leads to uncontrolled secretion of fluid and electrolytes into the lumen of the small intestine^40–42^. Assuming CTXA exerts its lethality in our yeast model through ADP ribosylation, therefore, blocking ADP ribosylation should lessen the lethality of cholera toxin. Keeping this in mind, we selected novobiocin, a known mono ADP ribosylation inhibitor^43^ and examined its effect on the reversal of toxin mediated toxicity in the recombinant yeast. We observed novobiocin rescued yeast growth similar to apomorphine (**Figure 3A and Figure 3B**).

**Figure 3.**
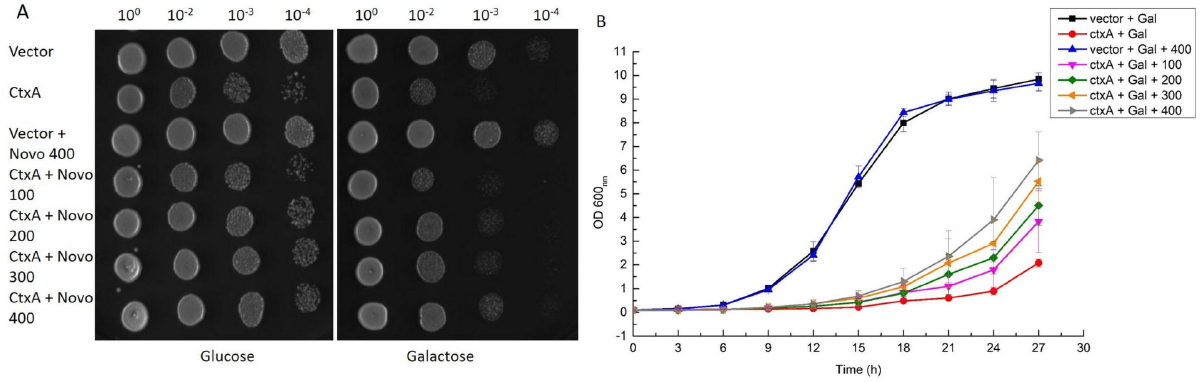
Effect of novobiocin on cholera toxicity in yeast: Secondary cultures of BY4741/ctxA-HO and vector control were set up at starting OD_600_ 0.05 in glucose and galactose media with different concentrations of novobiocin and were monitored for 27 hrs. OD_600_ was checked every 3 hrs. The data shown here is collected from 3 independent experiments (**A)**. At the end point of liquid growth assay, the cultures were serially diluted (10^0^, 10^1^, 10^2^, 10^3^) and were spotted on glucose and galactose plates. Pictures were taken after 60-70 hrs **(B)**. Liquid growth assay.

### A combination of apomorphine and novobiocin restricted CT mediated cellular toxicity and exhibited better survival of intestinal cells

Cholera toxin (CT) causes morphological changes of intestinal epithelial cells^34^, therefore, we desired to investigate whether CT mediated morphological changes and cellular death can be restricted by the exogenous administration of apomorphine and novobiocin. Towards this end, we treated HT29, a well characterised intestinal epithelial cell with cholera toxin (Sigma) with various concentrations of cholera toxin and observed a dose dependent morphological changes of HT29 cells (**Figure 4A**). Survival of intestinal cell was reduced approximately to 50-55 % in the presence of cholera toxin after 24 hr of treatment as measured by flow cytometry analysis (**Figure 4B**). To examine whether apomorphine and novobiocin can reverse cellular death caused by cholera toxin, various concentrations of drugs were added simultaneously with different concentrations cholera toxin on HT29 cells. After stipulated period of incubation, cellular morphology was microscopically examined. We observed both apomorphine and novobiocin effectively blocked CT mediated morphological changes and maintained wild type morphology of HT29 cells (**Figure 4A**). To gain further insight on the viability status of epithelial cells, a FACS based live-dead assay was performed. We observed significant rescue of toxin treated cells in the presence of drugs. Importantly, combined administration of both drugs strongly mitigated CT induced toxicity and promoted cellular survival (**Figure 4B**). Control sets containing drugs either alone or together did not affect the health of intestinal cells at the highest concentration (**Supplementary Figure 1**).

**Figure 4.**
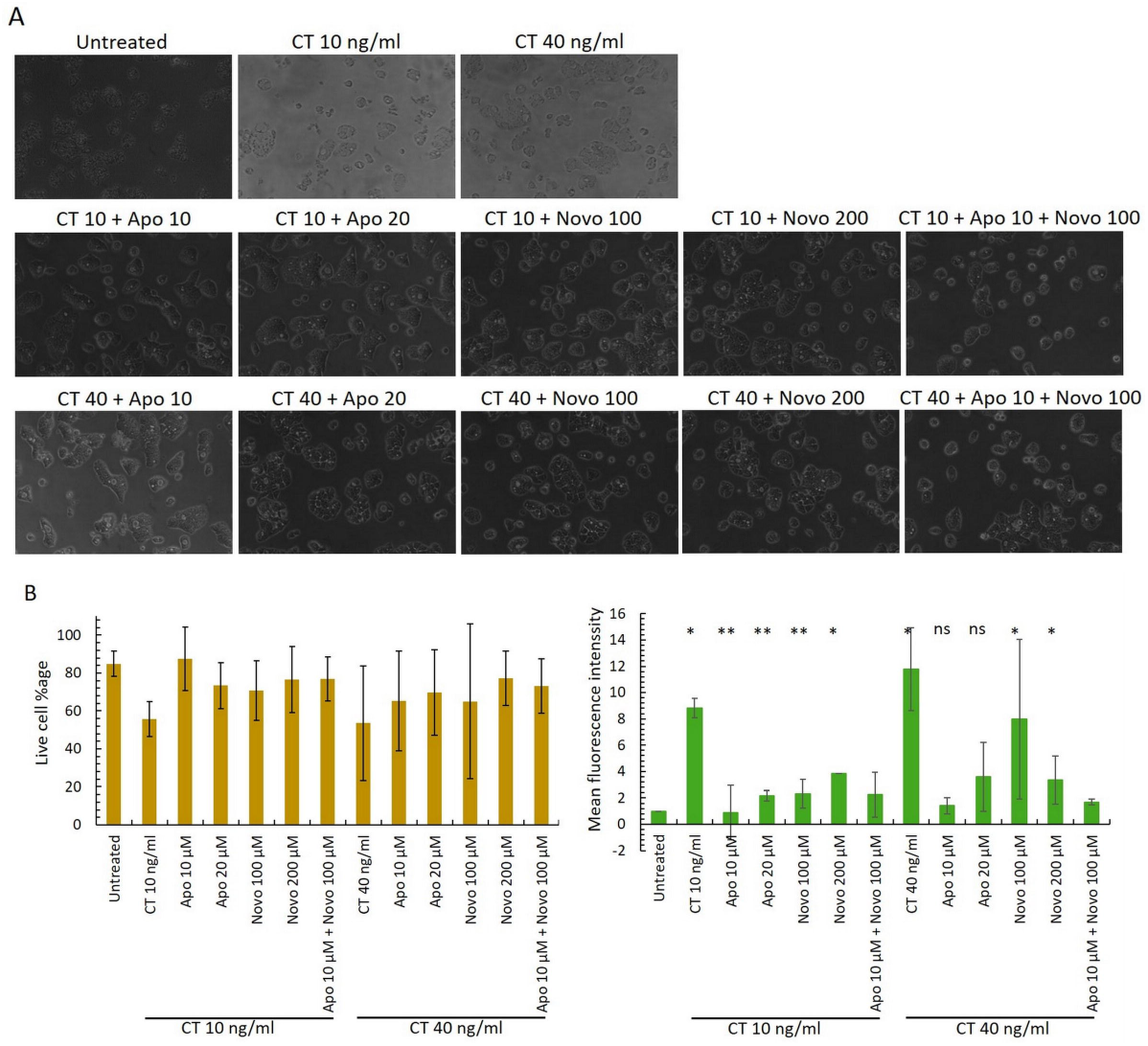
Effect of apomorphine HCl and novobiocin on CT toxicity on HT29: **A.** 0.2 million HT29 cells were seeded in 24 well plates. At 60% confluence, cells were washed 2X with DPBS. cholera toxin and drugs treatment was given for 6 hrs. Images were taken at 20X magnification under a light microscope. **B. Flow cytometry analysis:** Left panel shows the comparison between cell viability after 24 hrs of toxin and drug treatments and the right panel shows the mean fluorescence intensity of the same samples. The data shown here is collected from 2 independent experiments. Statistical analysis was performed using student’s T test (* P<0.05, ** P<0.01, NS-not significant).

## Discussion

Cholera is still one of the major killer diseases in the developing and war-torn countries. As summarized in the introductory section, there exists a demand towards the development of newer strategies balancing therapeutic safety and efficacy to control the disease in a better way. Keeping this in mind, we embarked on the present program to construct a yeast model of cholera toxin and subject this model to screen a chemical library. We choose yeast model primarily because of our prior engagement with yeast model of type III effectors^28,30,31^ and yeast models have been demonstrated highly effective and instrumental in screening small molecule libraries^32,33^. The library “Spectrum collection” from Microsource Discovery systems Inc. used in our study also exploited by other groups extensively for the discovery of many novel and repurposing drug molecules some of which are now under clinical trials^44^. Keeping view on various successful repurposing drug leads obtained from this library, we started our screening program with this library. Drug repurposing has several advantages over conventional drug discovery programmes in terms of safety and product development.

Various different methods are used to find out new applications of drug molecules that are different from its original course of action includes *in silico* techniques, literature survey and screening drug libraries. Armed with these strategies, our combined effort on literature survey and screening small molecule library eventually aid in the discovery two molecules namely novobiocin and apomorphine which are inhibiting cholera toxin mediated cellular lethality.

Novobiocin is an aminocoumarine antibiotic produced by actinomycete *Streptomyces niveus* and used to treat MRSA (methicillin resistant *Staphylococcus aureus*) infection. Novobiocin targets DNA gyrase which is a bacterial enzyme that functions as a catalyst for ATP dependent negative supercoiling of closed circular double stranded DNA^45,46^. The molecule also blocks HSP90 (heat shock protein of 90 kDa) chaperone protein in eukaryotic cells. This newly identified function of novobiocin is being explored as a possible anticancer agent^47^. Novobiocin also is an arginine dependent ADP-ribosyl transferase (ART) inhibitor compound. It has been widely used in studies of ARTs to determine their biological functions. This function of novobiocin was exploited in our present study. We observed that novobiocin is able to counter cholera toxin mediated toxicity in our experimental model systems. Considering its function as an ART inhibitor, it is conceivable that novobiocin may interfere toxin mediated massive production of cAMP, thereby, mitigating toxin mediated cellular lethality.

Apomorphine HCl is clinically used to treat Parkinson’s disease. The molecule is also explored as a possible treatment for Alzheimers disease^48,49^. The molecule is a known dopamine D2 receptor agonist. There are five types of dopamine receptors (D1 to D5), and each receptor has a different function. These five receptors are further divided into two subcategories: D1 and D5 are associated with Gs alpha subunit of G protein coupled receptor (GPCR) and D2, D3, D4 are coupled with Gi subunit of GPCR. Dopamine function in brain depends on the receptor that it binds to e.g., if it binds to D1R, it will lead to activation of adenylate cyclase through Gs alpha subunit and if it binds to D2R it will inhibit the enzyme through Gi subunit of GPCR^50^.

Being a D2 receptor agonist, we surmised that binding of apomorphine to its cognate receptor results inhibition of adenylate cyclase, thereby, perturbing cholera toxin mediated production of cAMP. Based on our observation, a molecular road map of drug and toxin interaction is shown as schematic diagram (**Figure 5**). Additional studies are necessary to address the issues.

**Figure 5:**
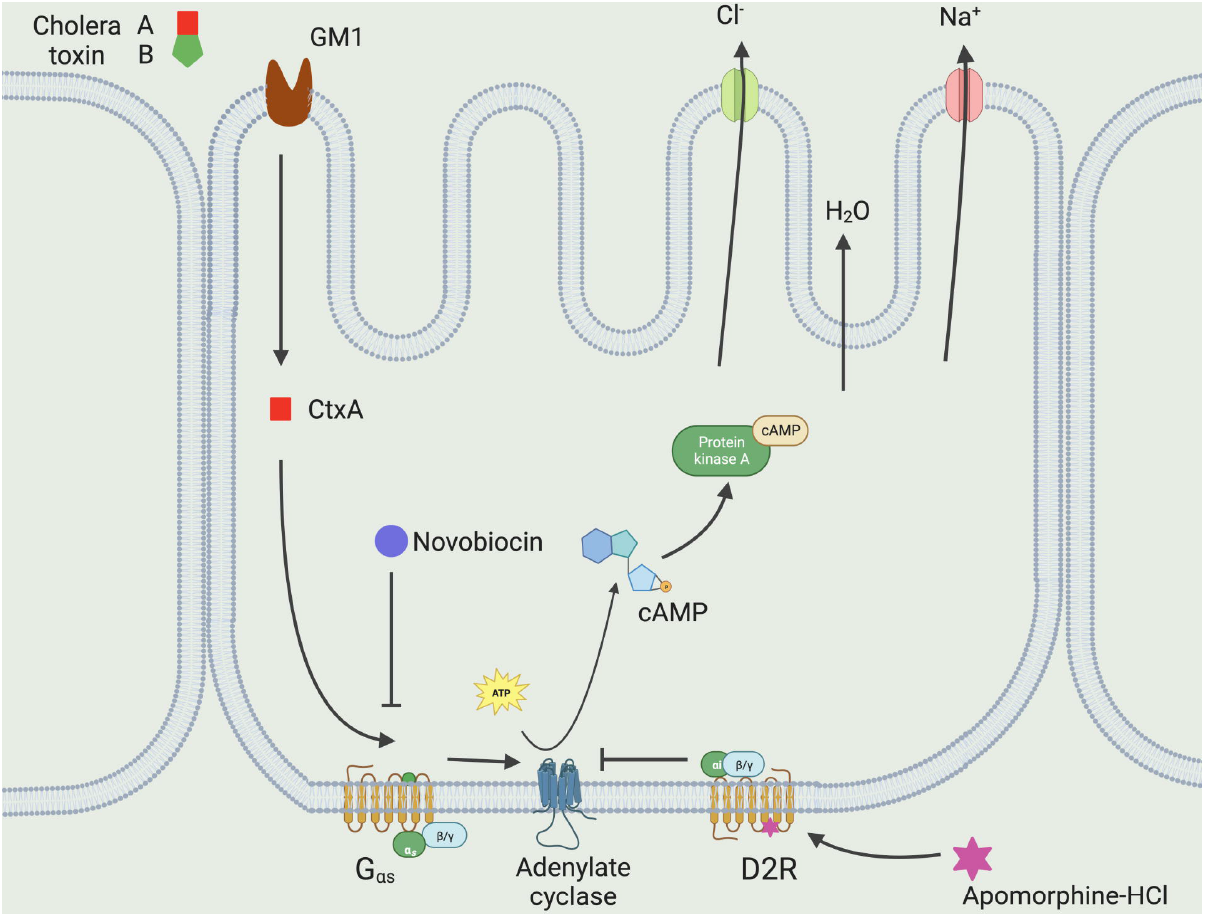
Predicted mechanism of action of apomorphine HCl and novobiocin against cholera toxin action. Schematic was adapted from Rahman et al., 2018^50^

## Materials and Methods

### Yeast strains and culture conditions

The *S. cerevisiae* strain BY4741 used in this study was grown in YPD (1% yeast extract, 2% peptone, 2% glucose) broth or agar plates (2%) at 30°C. Yeast strains harbouring plasmids were cultured in appropriate selective SC (synthetic complete) liquid media (0.67% Yeast nitrogen base without amino acids, 2% glucose, addition appropriate amino acids and nucleotide bases) or agar plates (2%). For 100 ml media, the concentration of amino acids and bases is used as follow: methionine 4 mg, histidine 4 mg, leucine 6 mg, tryptophan 4 mg, adenine 4 mg and uracil 4 mg. In case of SC^Raff^ and SC^Gal^ media, glucose is replaced with raffinose and galactose respectively. Yeast strains transformed with KanMX4 selection marker were selected by plating them on YPD agar plates containing G418 (200 μg/ml).

### Cell line and culture conditions

Human intestinal epithelial cell line (HT29) were obtained from Dr. Ravi Mishra lab (ATCC HTB-38) and were cultured in RPMI 1640 media obtained from Gibco, Invitrogen with 10% heat inactivated FBS and 1X antibiotic-antimycotic from Gibco. Cell cultures were incubated at 37°C, 5% CO_2_.

### Cloning

Gene encoding cholera toxin A subunit without signal sequence (highlighted in grey) was cloned under a GAL10 promoter into pESC-Leu, pGML10 vectors and HO-locus of the chromosomal DNA of yeast strain of BY4741. The primers used for the cloning of *ctxA* are listed along with the sequences in table 1.

>CTXA

ATGGTAAAGATAATATTTGTGTTTTTTATTTTCTTATCATCATTTTCATATGCAAATGATGATAAGTTATATCGGGCAGATTCTAGACCTCCTGATGAAATAAAGCAGTCAGGTGGTCTTATGCCAAGAGGACAGAGTGAGTACTTTGACCGAGGTACTCAAATGAATATCAACCTTTATGATCATGCAAGAGGAACTCAGACGGGATTTGTTAGGCACGATGATGGATATGTTTCCACCTCAATTAGTTTGAGAAGTGCCCACTTAGTGGGTCAAACTATATTGTCTGGTCATTCTACTTATTATATATATGTTATAGCCACTGCACCCAACATGTTTAACGTTAATGATGTATTAGGGGCATACAGTCCTCATCCAGATGAACAAGAAGTTTCTGCTTTAGGTGGGATTCCATACTCCCAAATATATGGATGGTATCGAGTTCATTTTGGGGTGCTTGATGAACAATTACATCGTAATAGGGGCTACAGAGATAGATATTACAGTAACTTAGATATTGCTCCAGCAGCAGATGGTTATGGATTGGCAGGTTTCCCTCCGGAGCATAGAGCTTGGAGGGAAGAGCCGTGGATTCATCATGCACCGCCGGGTTGTGGGAATGCTCCAAGATCATCGATGAGTAATACTTGCGATGAAAAAA CCCAAAGTCTAGGTGTAAAATTCCTTGACGAATACCAATCTAAAGTTAAAAGACAAATATTTTCAGGCTATCAATCTGATATTGATACACATAATAGAATTAAGGATGAATTATGA

### Yeast growth assay

The desired plasmid constructs were transformed into *S. cerevisiae* strain BY4741. The recombinant strains were selected by plating on selective SC solid agar plates. Three colonies of each clone were inoculated into selective SC^raf^ media and grown overnight at 30°C. The overnight grown cultures were used to set up a secondary culture in a fresh media with a starting A600 of 0.1, the cultures were allowed to grow till mid log phase. The effect of cholera toxin on yeast was examined by spotting equal number of cells on SC and SC^Gal^ plates lacking corresponding auxotrophic markers to maintain a plasmid. Yeast growth was monitored for 60-70 hrs and images were documented. For liquid growth assay, the cultures grown in SC^raf^ media were diluted to A_600_ of 0.05 in 25 ml of induction media (galactose) with appropriate nutritional supplementation. A_600_ was measured for stipulated period.

### Small molecule library screening

Primary screening was done in 96 well plates using yeast clones BY4741/pGML10 and BY4741/*CtxA* no signal/pGML10. Spectrum collection small molecule library (MicroSource Discovery Systems Inc.) was used to screen molecules against cholera toxin in yeast model. Overnight grown cultures were used to set up a secondary culture of BY4741/pGML10 and BY4741/*ctxA* no signal/pGML10 at A600 = 0.15 in YNB-Glucose-HMU media. At OD_600_ 0.3-0.4, cultures were induced with 2% galactose and were further incubated at 30°C for 4 hrs on shaking. After 4 hrs, ampicillin (50 μM) was added to the cultures to avoid bacterial contamination, and cultures were distributed in 96 well plates (160 μl/well). 20 μM of compounds were also added in 96 well plates from small molecule library. Using multichannel pipette, cultures and compounds were mixed well and plated were incubated for 24 hrs at 30 °C on static. The effect of drug molecules on cholera toxicity was examined by spotting the cultures on SC and SC^Gal^ plates using prong replicator. Yeast growth was monitored for 60-70 hrs and images were documented to examine growth restoration analysis.

### Cell line morphology study

HT29 cells were cultured in 12 well plate to 60% confluence. Media was discarded and cells were washed 2X with 1X DPBS and fresh media was added to the wells. Cholera toxin obtained from sigma (10ng/ml and 40ng/ml) and drug molecules (apomorphine-10 μM, 20 μM and novobiocin-100 μM, 200 μM, Apo 10 μM + Novo 100 μM) were added to the wells and mixed properly by gentle shaking. Plates were kept at 37°C, 5% CO_2_ for incubation. After 6 hrs, pictures were taken under light microscope at 20X resolution.

### Sample preparation for flow cytometry

HT29 cells were grown to 50% confluence in 12 well plate and were treated with CT (10 ng/ml and 40 ng/ml) and drug molecules (apomorphine-10 μM, 20 μM and novobiocin-100 μM, 200 μM, Apo 10 μM+ Novo 100 μM). After 24 hrs of treatment, media was collected in MCTs and was centrifuged at 850g for 10 mins to collect dead cells floating in the media and adhered cells were treated with trypsin in the wells. Supernatant was discarded from MCTs and trypsinized cells were collected in the same tubes and were centrifuged again at 850g for 5 mins. Positive controls were prepared by treating cells at 94°C for 3 mins. Cells were kept on ice for further treatment. 200 μl, propridium iodide stain (25 μg/ml) and 0.3 μl LIVE/DEAD™ fixable green dead cell stain from Thermofisher scientific was added to the culture pellets, mixed properly and were kept on ice for 10 mins. After staining, cells were washed with 1X DPBS and were resuspended in 500 μl 1X DPBS. Samples were sorted immediately with BD FACSVerse cell analyzer using 2 different excitation emission spectrums of wavelengths 488 nm and 566 nm. Data were analyzed using FlowJo v10 software.

## Acknowledgements

We gratefully acknowledge Dr. Deepak Sharma, Dr. Ravi Mishra and Dr. Ashwani Kumar for providing small molecule library and cell culture facility. This work was partly supported by grants from CSIR-IMTECH (OLP 151), CSIR-MLP39 and Science and Engineering Research Board (CRG/2018/000297/SERB-GAP/0185). Sonali E Bhalerao and Himanshu Sen, acknowledge Council of Scientific and Industrial Research (CSIR) for fellowships. Sonali E Bhalerao also acknowledges MLP 0039.

## Author contributions

SRC conceived the idea. SRC and SB designed the experiments. SB carried out all experiments, HS repeated experiments. SB, HS and SRC reviewed all data. SRC wrote the manuscript. All the authors gave editorial input.

---

**Supplementary Fig 1-Effect of apomorphine HCl and novobiocin on HT29: A.** 0.2 million HT29 cells were seeded per well. At 60% confluence, cells were washed 2X with DPBS. Drugs treatment was given for 6 hrs. Images were taken at 20X magnification under a light microscope. **B. Flow cytometry analysis:** Comparison between cell viability after 24 hrs of drugs treatment. The data shown here is collected from 2 independent experiments.

