## Supplemental tables for "Combined administration of novobiocin and apomorphine mitigates cholera toxin mediated cellular toxicity"

**Table 1: Primers used in the study**

| name | SEQUENCE |
| --- | --- |
| XhoI/*ctxA* (pESC-Leu) (FP) | 5’ - CCGCTCGAGATGAATGATGATAAGTTATATC - 3’ |
| NheI/*ctxA*  (pESC-Leu) (RP) | 5’ - CTAGCTAGCTCATAATTCATCCTTAATTC- 3’ |
| BamHI/ *ctxA* (pGML10) (FP) | 5’- CGCGGATCCATGAATGATGATAAGTTATATC - 3’ |
| EcoRI /*ctxA* (pGML10) (RP) | 5’- CCGGAATTCTCATAACTCATCCTTAATTCT -3’ |
| BamHI/ *ctxA* (HO-Locus) (FP) | 5’ - CGCGGATCCATGAATGATGATAAGTTATATC -3’ |
| SmaI /*ctxA* (HO-Locus) (RP) | 5’- TCCCCCGGGTCATAACTCATCCTTAATTCT - 3’ |

KEY RESOURCES TABLE

| REAGENT or RESOURCE | SOURCE | IDENTIFIER |
| --- | --- | --- |
| Bacterial and yeast strains and cell lines | | |
| DH5α | Novagen | 70181 |
| BY4741 | Bankapalli *et al., 2015, Bankapalli et al., 2017* | N/A |
| HT29 | Gift from Dr Ravi Mishra | ATCC HTB-38 |
| Plasmids | | |
| pESC-Leu | Agilent technologies | #217452 |
| pGML10 | RIKEN | RDB01957 |
| HO-pGal-polyKanMX4-HO | Addgene | #51664 |
| Chemicals and reagents | | |
| Yeast extract | BD Difco | 212750 |
| Tryptone | BD Difco | 211705 |
| NaCl | Sigma Aldrich | S9889 |
| Bacto agar | BD Difco | 214010 |
| Yeast nitrogen base | BD Difco | 291940 |
| Glucose | Sigma Aldrich | G8270 |
| Galactose | Sigma Aldrich | G0750 |
| Raffinose | Sigma Aldrich | R0250 |
| Histidine | Sigma Aldrich | H8000 |
| Methionine | Sigma Aldrich | M9625 |
| Uracil | Sigma Aldrich | U0750 |
| Tryptophan | Sigma Aldrich | 51145 |
| Adenine | Sigma Aldrich | A9126 |
| G418 | Calbiochem | 108321-42-2 |
| Ampicillin | Sigma Aldrich | A9393 |
| RPMI-1640 | Gibco, Invitrogen | 11875085 |
| Heat inactivated FBS | Gibco, Invitrogen | 10082147 |
| Antibiotic-antimycotic | Gibco, Invitrogen | 15240062 |
| 0.25% trypsin with EDTA | Gibco, Invitrogen | 25200072 |
| DPBS | Gibco, Invitrogen | 14200075 |
| Propridium iodide | Sigma Aldrich | P4170 |
| Spectrum collection small molecule library | Gift from Dr Deepak Sharma | MicroSource Discovery Systems Inc., Gaylordsville |
| Cholera toxin | Sigma Aldrich | C8052 |
| Apomorphine-HCl | Abcam | ab269887 |
| Novobiocin sodium salt | Sigma Aldrich | N1628 |
| Critical commercial assays | | |
| Pierce BCA Protein Assay | Thermofisher scientific | #23227 |
| LIVE/DEAD™ Fixable Green Dead Cell Stain Kit | Thermofisher scientific | L34970 |
