## Supplementary figures and images for "Combined administration of novobiocin and apomorphine mitigates cholera toxin mediated cellular toxicity"

### Supplemental Fig

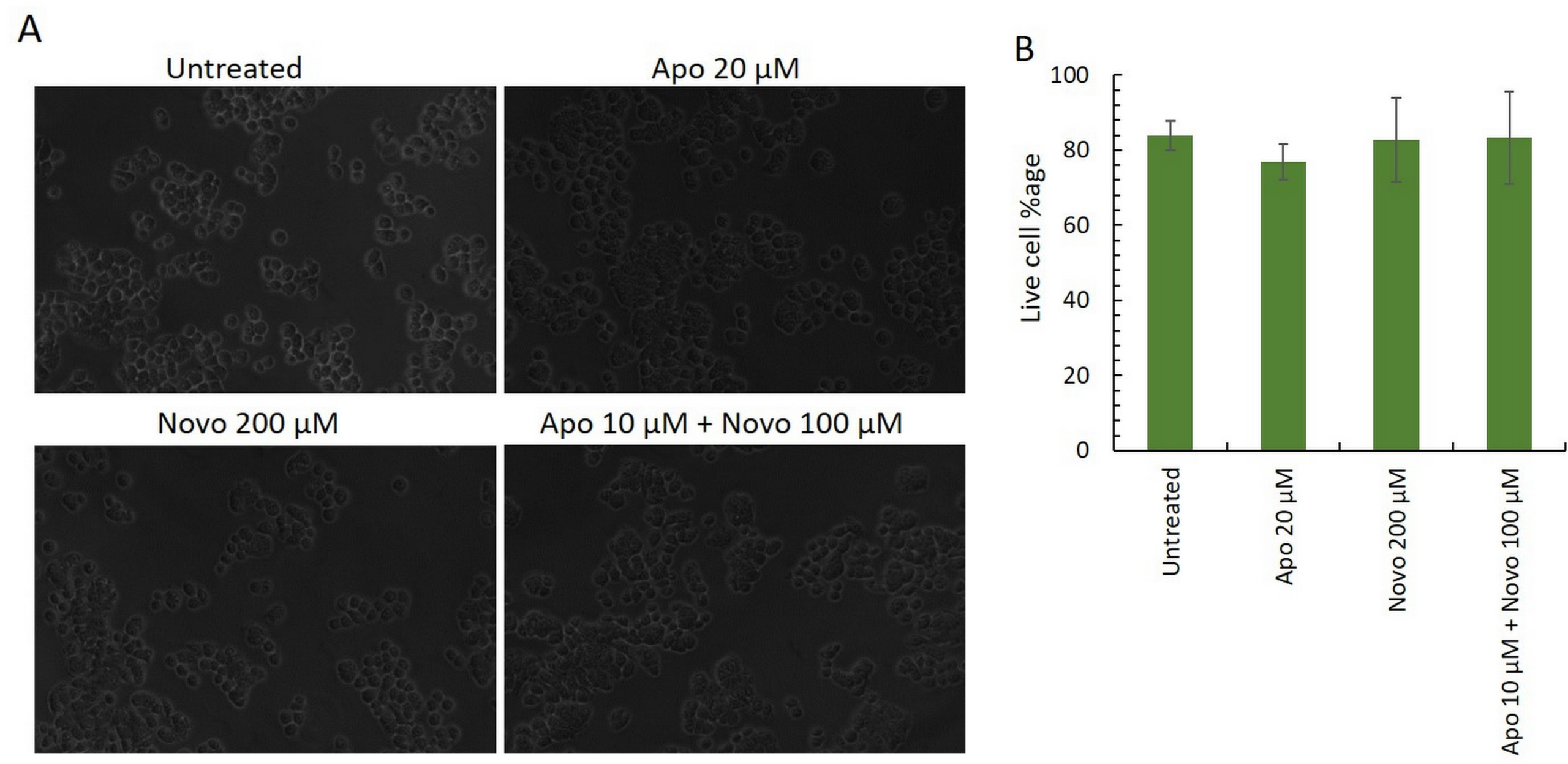
